# Absolute quantification of photoreceptor outer segment proteins

**DOI:** 10.1101/2023.01.19.524794

**Authors:** Nikolai P. Skiba, Tylor R. Lewis, William J. Spencer, Carson M. Castillo, Andrej Shevchenko, Vadim Y. Arshavsky

**Author notes:** Department of Ophthalmology and Visual Sciences, SUNY Upstate Medical University, Syracuse, NY 13210.

## Abstract

Photoreceptor cells generate neuronal signals in response to capturing light. This process, called phototransduction, takes place in a highly specialized outer segment organelle. There are significant discrepancies in the reported amounts of many proteins supporting this process, particularly those of low abundance, which limits our understanding of their molecular organization and function. In this study, we used quantitative mass spectrometry to simultaneously determine the abundances of twenty key structural and functional proteins residing in mouse rod outer segments. We computed the absolute number of molecules of each protein residing within an individual outer segment and the molar ratio amongst all twenty proteins. The molar ratios of proteins comprising three well-characterized constitutive complexes in outer segments differed from the established subunit stoichiometries of these complexes by less than 7%, highlighting the exceptional precision of our quantification. Overall, this study resolves multiple existing discrepancies regarding the outer segment abundances of these proteins, thereby advancing our understanding of how the phototransduction pathway functions as a single, well-coordinated molecular ensemble.

## Introduction

Rod and cone photoreceptors are retinal neurons which generate electrical signals in response to capturing light. This process, called phototransduction, takes place in the outer segment organelle that contains a stack of membranous “discs” to facilitate efficient light capture. The molecular composition of the outer segment is rather unique and includes a number of functional and structural proteins specialized in supporting phototransduction. Over the past several decades, most of these proteins have been characterized in great detail, which makes phototransduction one of the most robust systems to study the general principles of GPCR signaling and one of the best understood signaling cascades overall [1, 2]. Not surprisingly, the phototransduction cascade has been the subject of extensive computational modeling and is viewed as a powerful quantitative system for appreciating the coordination of individual proteins within a complex network (e.g., [3-7]). However, a complete quantitative understanding of phototransduction is still limited by insufficient information on the precise outer segment contents of many phototransduction proteins, particularly those of lower abundances. The goal of this study was to address this knowledge gap.

Nearly all existing information on protein amounts in the outer segment was obtained from either semi-quantitative measurements of individual or small sets of proteins (e.g. [8, 9]) or rough estimates from large proteomic datasets based on relative representations of peptide spectral counts [10]. Accordingly, there is a great disparity across published abundances for many of these proteins (e.g. [11] vs. [12] or [13] vs. [14]; see also **Table 1** below). A more accurate mass spectrometry-based approach for absolute protein quantification involves the use of commercially synthesized isotope labeled peptide standards corresponding to tryptic fragments of a protein of interest. To our knowledge, this technique has only been applied to quantifying a handful of outer segment proteins in a single study [15]. The limited use of this approach is due, at least in part, to the high costs of labeled peptide standards, potential issues with the stability and solubility of these peptides and the fact that concentrations of many commercial peptide standards are not easy to independently verify in a quantitatively consistent manner [16].

**Table 1.** Summary of protein quantification in dark-adapted mouse rod outer segments

| Protein | Ratio to rhodopsin | Coefficient of variation, % | Molecules per rod outer segment | Molecules per disc | Reported ratio to rhodopsin |
| --- | --- | --- | --- | --- | --- |
| Rhodopsin | - | - | 60,000,000<sup>&</sup> | 75,000 | - |
| G$\alpha_t$ | 1:7.8 | 2.9 | 7,690,000 | 9,600 | 1:8 [42]; 1:8 [48]; 1:12 [26]; 1:8* [69] |
| G$\beta_1$ | 1:7.9 | 2.2 | 7,590,000 | 9,500 | 1:8 [42]; 1:6 [48], 1:12 [26] 1:10* [69] |
| PDE6α | 1:132 | 6.6 | 455,000 | 570 | 1:120 [42]; 1:40-170 [43]; 1:65-104 [44]; 1:310 [45]; 1:200* [69], 1:270* [77]; 1:330* [78] |
| PDE6β | 1:141 | 7.7 | 426,000 | 530 | |
| GRK1 | 1:201 | 8.9 | 299,000 | 370 | 1:360 [13]; 1:40 [14] |
| Arrestin | 1:18 | 12 | 3,340,000 | 4,200 | 1:7-10 [51]; ≤1:18 [52]; 1:33* [69] |
| Recoverin | 1:111 | 27 | 541,000 | 670 | 1:104 [50]; 1:174* [79] |
| RGS9 | 1:461 | 5.8 | 130,000 | 170 | 1:1,640 [11]; 1:610-1:910 [44]; 1:269 [12] |
| G$\beta_5$ | 1:469 | 6.2 | 128,000 | 160 | |
| R9AP | 1:474 | 9.6 | 127,000 | 160 |  |
| CNG$\alpha_1$<sup>#</sup> | 1:323 | 6.2 | 185,000 | 230 | ~1:500 for the tetrameric channel [46] |
| CNG$\beta_1$<sup>#</sup> | 1:988 | 9.2 | 60,700 | 76 | |
| NCKX1 | 1:472 | 7.4 | 127,000 | 160 | 1:800 [54] |
| Guanylyl cyclase 1 | 1:526 | 8.2 | 114,000 | 140 | 1:440 [59]; 1:260 [8]; 1:1,400 [9] |
| Guanylyl cyclase 2 | 1:3,490 | 7.2 | 17,200 | 21 | 1:6,600 [60]; 1:5,800 [9] |
| Peripherin-2 | 1:19 | 8.9 | 3,160,000 | 3,900 | 1:90 for the peripherin-2/ROM1 tetramer [66]; calculated as 1:25 for the tetramer [67] |
| ROM1 | 1:39 | 6.0 | 1,540,000 | 1,900 |  |
| ABCA4 | 1:794 | 5.2 | 75,600 | 95 | 1:302 [15], 1:120 [80], 1:167* [69] |
| ATP8A2 | 1:13,150 | 11 | 4,560 | 8 | <0.1% of total outer segment protein [70] |

<sup>&</sup>This value for rhodopsin is averaged from [33, 34].
<sup>\*</sup>These values were determined for frog rather than mammalian outer segments.
<sup>#</sup>The molar ratio to rhodopsin is calculated to be 1:966 for the tetrameric CNG channel.

These limitations have been overcome with the introduction of a strategy for absolute protein quantification using the digest of a single recombinant protein built as a concatemer of tryptic peptide standards [17, 18]. This cost-efficient approach does not require synthesizing individual peptide standards and allows to quantify multiple proteins in a single analysis. In this study, we employed a variation of this approach, termed “MS Western” [19], in which sequences of several tryptic peptides from each protein of interest are cloned in tandem as a part of a single DNA construct that also encodes several tryptic peptides from BSA. This chimeric protein is expressed in an *E. coli* strain auxotrophic for arginine and lysine grown in a medium containing heavy isotope labeled lysine and arginine. A tryptic digest of this heavy isotope labeled protein produces an equimolar set of its constituent peptides, including those from BSA. The chimeric protein is digested alongside unlabeled BSA of known amount and the sample of interest. The resulting peptide mixture is analyzed by LC-MS/MS and the absolute amounts of heavy isotope labeled BSA peptides from the chimera are determined by comparison to the corresponding peptides from unlabeled BSA whose precise amount is known. Because all other labeled peptides in the chimeric protein digest are equimolar to labeled BSA peptides, these labeled peptides can be used as standards for determining absolute (molar) amounts of each corresponding protein in the unlabeled sample of interest. This methodology has been successfully applied to the quantification of core histones in zebrafish embryogenesis [19], large sets of proteins in the *Drosophila* eye [20, 21] and central carbon metabolic enzymes in *C. elegans* [22, 23].

Using this approach, we determined the absolute abundances of twenty proteins supporting the function and structure of the mouse rod outer segment. The precision of the method was assessed by measuring the molar ratios amongst individual protein subunits from three constitutive protein complexes of well-established subunit stoichiometry. Strikingly, the deviation from known stoichiometric ratios did not exceed 7%, which highlights the exceptional precision of this method. This enabled us to establish a precise molar ratio amongst all analyzed proteins and to calculate their absolute amounts in each outer segment. These findings resolve multiple discrepancies in the literature regarding the abundances of some of these proteins and provide new quantitative information about the membrane flippase ATP8A2 whose abundance had not yet been analyzed.

## Experimental Procedures

### Animal husbandry

Animal maintenance and experiments were approved by the Duke Institutional Animal Care and Use Committee. We used C57BL/6J (Jackson Labs stock #000664) mice of both sexes between the ages of 1 and 6 months.

### Composition, expression and digestion of the chimeric protein

The polypeptide sequence of the chimeric protein, templated from [19], is illustrated schematically in **Supplementary Fig. 1**. It contained an N-terminal twin-strep-tag with a 3C protease cleavage site, concatenated sequences of 110 peptides (104 from proteins of interest and 6 from BSA; **Supplementary Table 1**), and a C-terminal His_6_-tag, although we did not need to use either tag for purification. DNA encoding this construct was custom synthesized and cloned into the pET11 vector at Genscript. It was transformed into competent ΔLys, ΔArg auxotrophic *E. coli* cells [24] generated and provided by Ronald Hay (University of Dundee). Transformed cells were grown in MDAG medium supplemented with ampicillin and ^15^N_4_,^13^C_6_-Arg and ^15^N_2_,^13^C_6_-Lys (Cambridge Isotope Laboratory). At OD600=0.6, cells were induced with 1 mM IPTG for 4 h. Cell lysates from 50 μl of cell culture were subjected to SDS-PAGE on a 10% gel. The protein band representing the chimeric protein was cut from the gel and cleaved using a standard in-gel trypsin digestion protocol [25]. The incorporation of ^15^N_4_,^13^C_6_-Arginine and ^15^N_2_,^13^C_6_-Lysine was 99.5% as judged by a ratio between the intensities of unlabeled and labeled peptides in a protein expressed in heavy isotope containing medium.

### Preparation and digestion of mouse rod outer segments

Mice were dark-adapted overnight and osmotically intact rod outer segments were isolated under dim red light following a protocol adapted from [26] with modifications described in [27]. Rhodopsin concentration in the resulting preparation was measured by differential spectroscopy at 500 nm as described in [28]. Samples containing 20-30 pmol rhodopsin were mixed with 100-250 fmol BSA and cleaved with 1 μg trypsin/LysC mix (Promega) using the SP3 beads protocol described in [29].

### Mass spectrometry

The combined digest of outer segments and BSA was mixed with 100-250 fmol of the chimeric protein digest and injected into a Symmetry C18 trap column (5 μm, 180 μm×20 mm) (Waters) in 1% acetonitrile in water at 5 μl/min. The analytical separation was subsequently performed using an HSS T3 column (1.8 μm, 75 μm×200 mm) (Waters) over 90 min at a flow rate of 0.3 μl/min at 50°C. The 5-30% mobile phase B gradient was used, where phase A was 0.1% formic acid in water and phase B was 0.1% formic acid in acetonitrile. Peptides separated by LC were introduced into the Q Exactive HF tandem mass spectrometer (Thermo Fisher Scientific) using positive electrospray ionization at 2 kV and ion transfer capillary temperature of 275°C. MS1 spectra were acquired in data-dependent acquisition (DDA) mode with the target mass resolution of 60,000 (*m/z* 200; FWHM); *m/z* range of 350-1350; target AGC value of 1×10^6^; RF-lens set at 30%; and maximum injection time of 50 ms. Advanced peak detection and internal calibration (EIC) were enabled. Peptide precursor ions were selected for MS/MS using charge state filtering (*z* = 2 to 5); monoisotopic peak detection and dynamic exclusion time of 20 sec with mass tolerance of 10 ppm. HCD FT MS/MS was performed under normalized collision energy (NCE) of 30%; AGC target value of 5×10^4^; maximum injection time of 100 ms; isolation width of precursor ions of 1.6 Th. DDA was guided by the inclusion list containing *m/z* of doubly and triply charged precursors of peptides used in subsequent absolute quantification.

### Data processing

For absolute protein quantification, raw mass spectral data files (.raw) were imported into Progenesis QI for Proteomics 4.2 software (Nonlinear Dynamics) for duplicate runs alignment of each preparation and peak area calculations. Peptides were identified using Mascot version 2.5.1 (Matrix Science) for searching a mouse UniProt 2019 reviewed database containing 17,008 entrees and the BSA sequence. Mascot search parameters were: 10 ppm mass tolerance for precursor ions; 0.025 Da for fragments mass tolerance; no missed cleavage allowed; fixed modification: carbamidomethylation of cysteine; variable modifications were oxidized methionine and N/Q deamidation and ICAT Lys_8_, Arg_10_ for isotope labels. False discovery rate for protein identification was set to <1%. Only peptides with ion scores more than 25 were included in quantification. To account for variations in experimental conditions in individual LC-MS/MS runs, the integrated peak area for each identified peptide was corrected using normalization to all proteins by a Progenesis algorithm equalizing the total intensities for all peaks in each run.

### Experimental design and statistical rationale

We analyzed a total of six independently obtained rod outer segment preparations. Each sample was run in duplicate and corresponding labeled and unlabeled peptide pairs were identified by Mascot software using “SILAC K+8, R+10” variable modification. The integrated areas of peptide peaks representing XIC of full isotope cluster were averaged between duplicates. The absolute amounts of heavy isotope labeled standards from the digest of the chimeric protein were determined using reference BSA peptides, whose abundance was compared with corresponding peptides produced by tryptic cleavage of BSA standard with the exactly known concentration. Next, the amounts of unlabeled peptides representing proteins of interest were determined based on their ratios to the amounts of labeled peptide standards. The amounts of proteins were calculated by averaging the amounts of their constituent peptides, after applying the exclusion criterion described in Results. The amounts of low abundant proteins were corrected for the presence of an impurity of non-labeled amino acids in the heavy isotope labeled chimeric protein standard. This impurity, identified to represent 0.5% of the added chimeric protein was subtracted from the calculated amount of the target protein. The data for each protein were averaged across all six outer segment preparations. Coefficient of variation was calculated using Excel.

## Results

The goal of this study was to determine the absolute amounts of proteins that perform phototransduction or otherwise uniquely reside in the photoreceptor disc membranes where phototransduction takes place. We only included proteins that were documented to produce at least two unique tryptic peptides confidently identified by mass spectrometry in our previous studies [15, 27, 30, 31]. This allowed us to analyze 16 proteins directly engaged in phototransduction (entries 1-16 in **Supplementary Table 1**) and four additional disc-specific proteins (entries 17-20). The roles of these proteins in photoreceptor function are illustrated schematically in **Fig. 1**. The four remaining phototransduction proteins (PDE6γ, Gγ_1_, GCAP1 and GCAP2) and the disc-specific protein PRCD were excluded from this study, as each yield only one tryptic peptide suitable for quantification.

**Figure 1.**
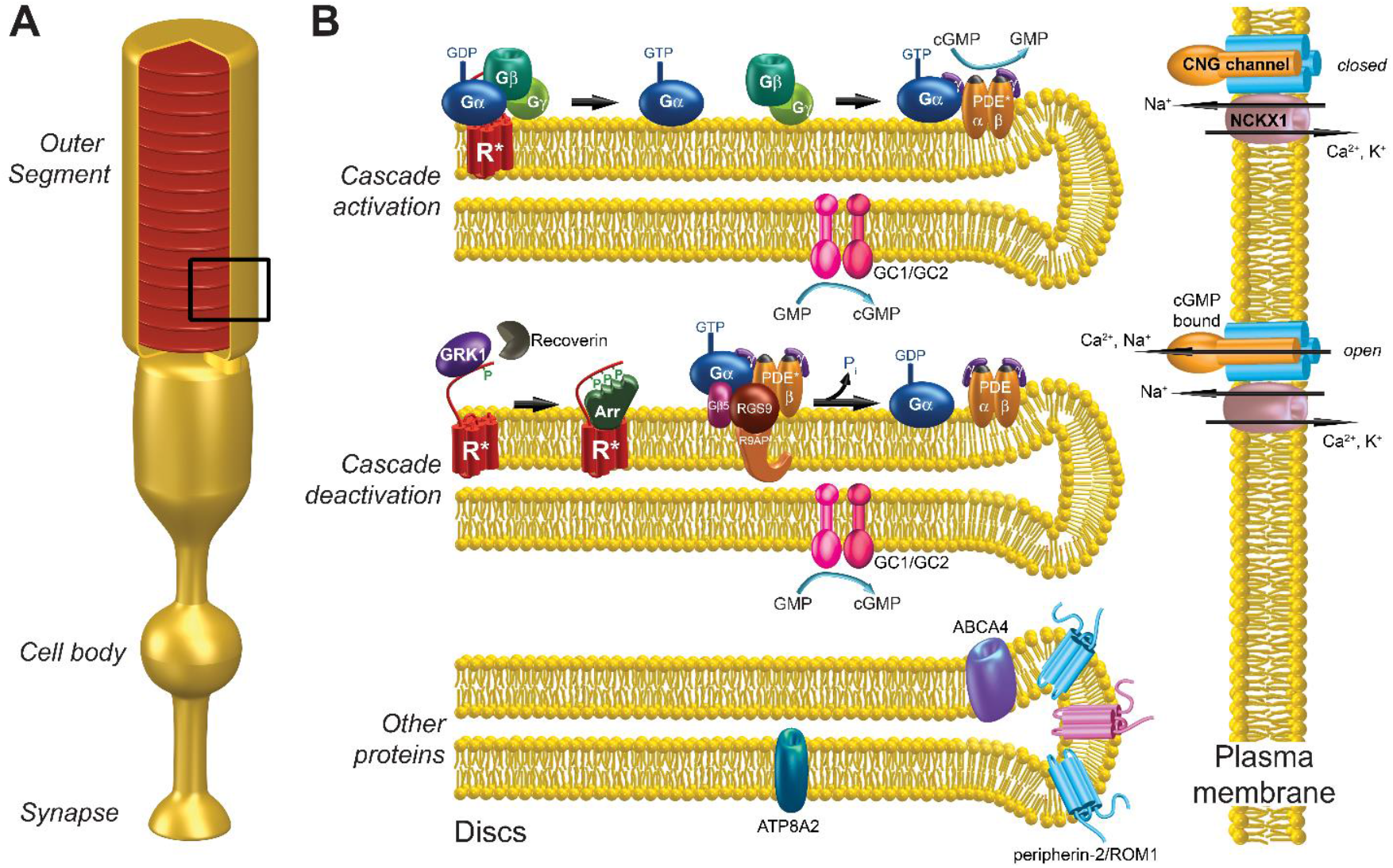
Cartoon illustrating localization and function of proteins analyzed in this study. **A**. Schematic of a rod photoreceptor cell. **B**. Zoomed-in view of the portion of the outer segment boxed in panel A. Illustrated at the *upper disc* is the activation of the phototransduction cascade, including photoexcited rhodopsin (R^*^), transducin (Gα and Gβγ) and PDE6 (PDEαβγ). Two guanylyl cyclase isoforms (GC1 and GC2) responsible for cGMP synthesis are shown on the bottom membrane leaflet. Illustrated at the *middle disc* are the reactions responsible for cascade deactivation. R^*^ is deactivated through phosphorylation by rhodopsin kinase (GRK1) and subsequent arrestin (Arr) binding; the activity of GRK1 is regulated by recoverin. Transducin deactivation occurs through GTP hydrolysis facilitated by the RGS9/Gβ5/R9AP GTPase activating complex; this returns PDE6 to its inactive state. The *plasma membrane* harbors the cGMP-gated (CNG) channel containing three α and one β subunits. This channel is open in the dark and closes upon cascade activation by light. The same membrane harbors the Na^+^/K^+^/Ca^2+^ exchanger (NCKX1). Illustrated at the *lower disc* are two tetraspanin proteins (peripherin-2 and ROM1) fortifying the disc rim and two lipid flippases (ABCA4 and ATP8A2). The drawing is modified with permission from [81].

For absolute quantification of most proteins of interest we were able to select 5 unique tryptic peptides (**Supplementary Table 1**) that have been consistently identified in our previous mass spectrometry studies [15, 27, 30, 31]. When possible, we excluded peptides containing methionine, tryptophan and cysteine due to a variable extent of their oxidation, as well as peptides with N-terminal aspartic acid due to a potentially decreased rate of trypsin cleavage at the Lys/Arg-Asp bond. The only protein that was represented by less than 5 peptides was the visual pigment rhodopsin, which yields only two tryptic peptides that are suitable for quantification.

To generate the chimeric protein standard, we cloned the sequences of these peptides in tandem as shown in **Supplementary Fig. 1**. Because rhodopsin is by far the most abundant protein in the outer segment, we included four copies of each of the two rhodopsin peptides in the chimera. This allowed us to keep the ratio between labeled and unlabeled rhodopsin peptides closer to the range for peptides representing other proteins and was accounted for in the final quantification. Behind the peptides representing outer segment proteins, we cloned 6 proteotypic peptides from BSA that were used in the original study describing the MS Western methodology [19]. The resulting construct also contained an N-terminal twin-strep-tag and 3C protease cleavage site and a C-terminal His_6_-tag that were parts of the original plasmid described in [19], but not utilized in the present study. This plasmid encoded a chimeric protein with a calculated molecular mass of 143 kDa. It was expressed in ΔLys, ΔArg auxotrophic *E. coli* [24] grown in the presence of ^13^C_6_, ^15^N_2_ lysine and ^13^C_6_ and ^15^N_4_ arginine. The band corresponding to this heavy isotope labeled chimeric protein was cut out of a Coomassie-stained SDS-PAGE gel (**Fig. 2A**) and trypsinized using a conventional in-gel digestion protocol [25].

**Figure 2.**
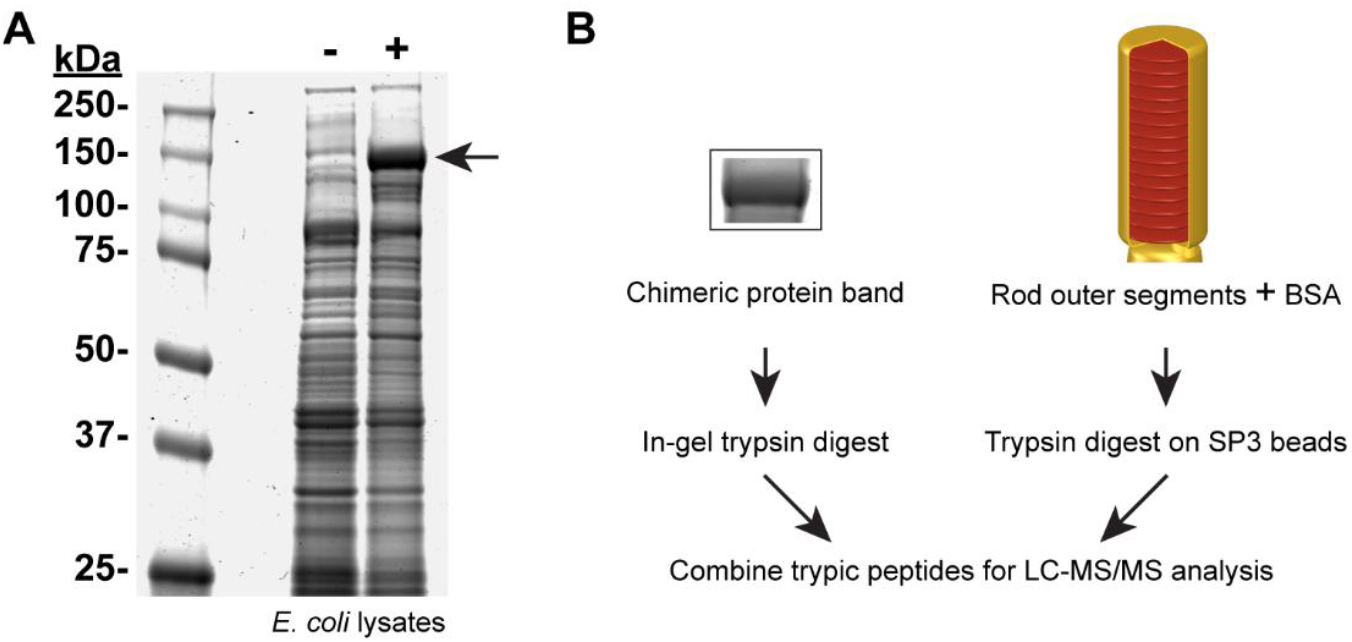
Experimental workflow for absolute quantification of outer segment proteins. **A**. Coomassie-stained SDS-PAGE gel of *E. coli* lysates (50 μl) from uninduced (-) and IPTG-induced (+) cells transformed with the DNA construct encoding the chimeric protein. Induction with IPTG caused a high level of expression of the chimeric protein migrating at the predicted molecular mass of ∼143 kDa (arrow). **B**. General experimental workflow for absolute quantification of outer segment proteins. A preparation of rod outer segments was combined with a known amount of BSA and digested with trypsin on SP3 beads. In parallel, the ∼143 kDa chimeric protein band was cut out from the Coomassie-stained SDS-PAGE gel and underwent in-gel trypsin digestion. The two digests were combined and subjected to LC-MS/MS analysis.

Our workflow for quantifying outer segment proteins is illustrated in **Fig. 2B**. We prepared a lysate of purified mouse rod outer segments, mixed it with BSA of known amount and trypsinized the mixture on SP3 beads as in [29]. Next, we supplemented this digest with the digest of the chimeric protein containing heavy isotope labeled peptide standards. We used a concentration of peptide standards corresponding to the middle of the concentration range for analyzed peptides. This workflow was modified from the original MS Western protocol, in which proteins in the sample, chimera and BSA were co-digested in-gel. The advantage of this modification was that the chimera was purified by SDS-PAGE and was not contaminated by truncated fragments and/or products of incomplete translation [32]. The concentration of The resulting sample was analyzed by LC-MS/MS in two technical repeats. The analysis was performed for six independently obtained outer segment preparations.

Whereas our peptide selection aimed to avoid common issues complicating analysis, such as oxidation or incomplete digestion, we could not predict whether any of these peptides would be differentially modified in (or unevenly recovered from) outer segment vs. *E. coli* lysates, thereby compromising their quantification. To determine whether any such peptides should be excluded from the final protein quantification, we used an approach described in [19], which was applied independently to the analysis of each of the six outer segment preparations. For each of the peptides representing a given protein of interest, we calculated the fraction of ion intensity that this peptide contributes to the total ion intensity of all peptides representing that protein. This was done separately for unlabeled and heavy isotope labeled peptide sets. If a pair of unlabeled and labeled peptides is equally digested and neither is modified, their relative abundances should be close to one another. This was the case for the majority of peptides, such as for all peptides representing RGS9 in the experiment illustrated in **Fig. 3A**. The peptides not satisfying this criterion, either in individual experiments or systematically across all experiments, were excluded from the analysis. The examples include one peptide representing Gβ_1_ (**Fig. 3B**) or two peptides representing peripherin-2 (**Fig. 3C**). The total number of systematically excluded peptides was six (they are labeled in red in **Supplementary Table 1**).

**Figure 3.**
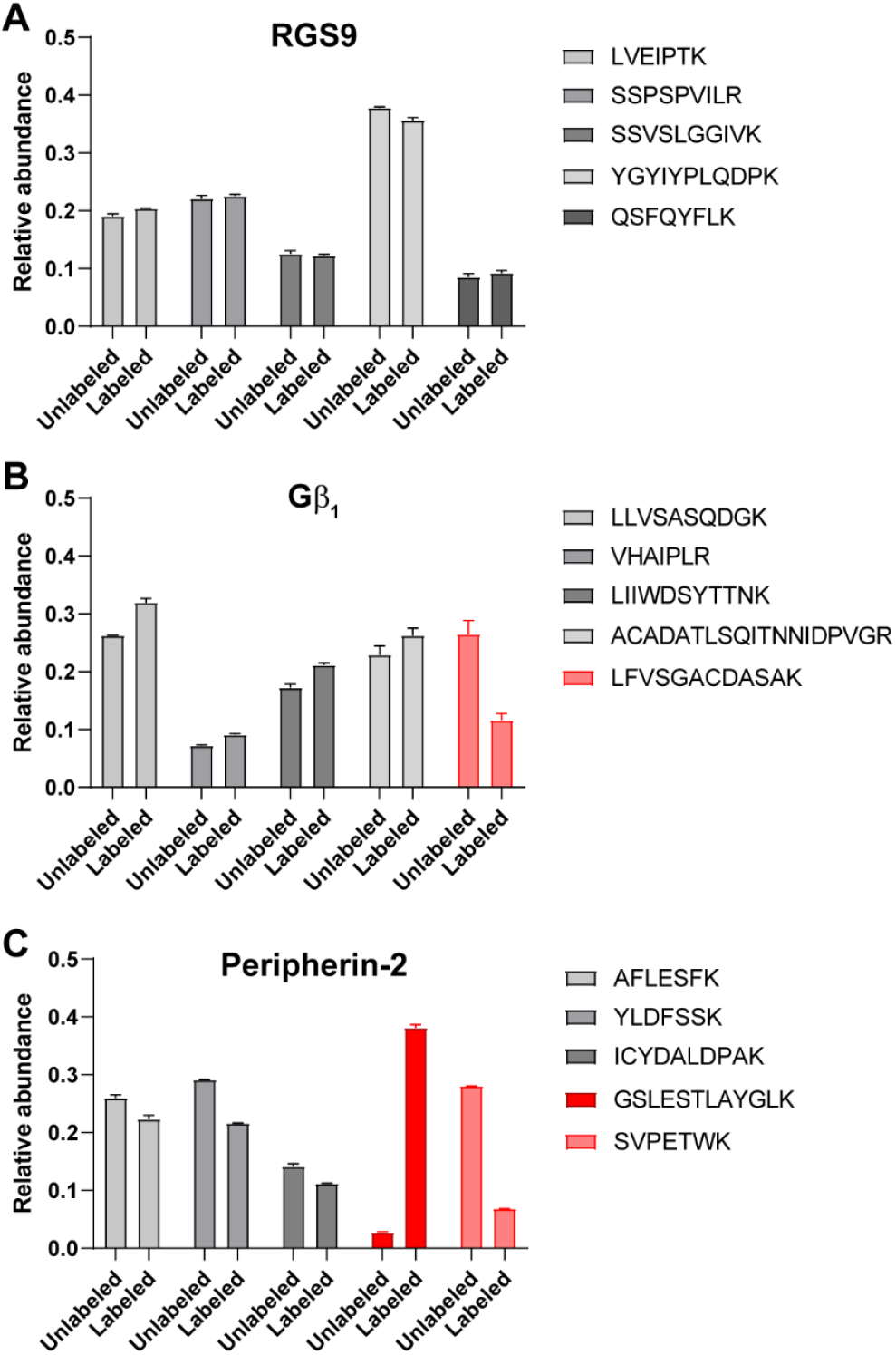
Peptide selection for final protein quantification. For each peptide representing a protein shown in each panel, the fraction of ion intensity that this peptide contributes to the total ion intensity of all peptides representing this protein was calculated separately for unlabeled and heavy isotope labeled peptide sets. **A**. Relative abundances of unlabeled and labeled peptides representing RGS9 in a single experiment with technical duplicates. In this case, the relative abundances of each unlabeled and the corresponding labeled peptide matched closely and satisfied the inclusion criterion. **B**. Relative abundances of unlabeled and labeled peptides representing Gβ_1_. The relative abundances of one unlabeled and labeled peptide pair (shown in red) were >2-fold different. The other four peptide pairs satisfied the inclusion criterion. **C**. Relative abundances of unlabeled and labeled peptides representing peripherin-2. The relative abundances of two unlabeled and labeled peptide pairs were >2-fold different. The other three peptide pairs satisfied the inclusion criterion.

We next quantified the absolute amounts of all 20 proteins of interest (**Supplementary Tables 1-6**). Because it is customary to describe the contents of outer segment proteins as molar ratios to rhodopsin, we calculated these ratios for all proteins (**Table 1**). We also calculated the number of molecules of each protein present in a single rod outer segment. This could be derived from the molar ratio of each protein to rhodopsin because the number of rhodopsin molecules per rod outer segment is known to be between 50 and 70 million [33, 34]. We used the average value of 60 million rhodopsin molecules per outer segment in this calculation. Finally, we estimated the abundance of each protein in the space within the outer segment occupied by a single photoreceptor disc, using an average value of ∼800 discs per mouse rod outer segment [35]. These estimates are summarized in **Table 1**, as well.

The analysis of phototransduction proteins provides a unique opportunity to assess the precision of absolute protein quantification using our methodology. Seven of the twenty proteins quantified in our study exist as constitutive multi-subunit protein complexes. The subunit stoichiometry of each of these complexes has been established by multiple independent approaches, including solved 3D structures, and can be used as a benchmark for validating the accuracy of protein quantification performed in our study. Furthermore, none of the proteins comprising these complexes are expressed in free form or as parts of other complexes. The first complex was the constitutive PDE6α/β dimer [36], for which our data deviated from the expected 1:1 stoichiometry by ∼6.8%. The second was the GTPase activating complex containing equimolar amounts of its three constituent subunits, RGS9, Gβ5 and R9AP [37]. Our measured amounts of each of these subunits were very close among themselves and differed from the expected 1:1:1 stoichiometry by only ∼2.8%. The third was the tetrameric cyclic nucleotide gated (CNG) channel that has a 3:1 molar ratio between CNGα1 and CNGβ1 [38-40]. We measured the ratio between these subunits to be 3.06 :1, which differs from the theoretical value by a remarkable 1%. These calculations illustrate the exceptional precision of the analysis employed in our study and suggest that this analysis provides the most accurate quantification of multiple outer segment proteins performed to date.

## Discussion

In this study, we report a precise molar ratio amongst the proteins comprising the phototransduction pathway and several other proteins supporting phototransduction in outer segment disc membranes. Remarkably, we were able to simultaneously quantify, in a single experiment, proteins whose outer segment abundances vary by three to four orders of magnitude.

### Methodological considerations

The approach employed in our study was modified from the original MS Western protein quantification method [19]. While the original protocol involved SDS-PAGE and in-gel digestion of the biological sample, chimeric protein and BSA standard, we only used SDS-PAGE and in-gel digestion for the chimeric protein. This was critical to ensure that the full-length chimeric protein was free from contaminations by its under-expressed or truncated fragments. In contrast, outer segment proteins and BSA were trypsinized on beads and, subsequently, combined with the digest of the chimera. Clearly, this modification did not affect the accuracy of quantification, yet it allowed us to use the same chimeric protein digest in multiple experiments. Another advance was to adjust the number of peptide copies in the chimera in accordance with the expected abundance of quantified proteins, in our case rhodopsin. This simplified the simultaneous quantification of multiple proteins of various abundances by easing the dynamic range requirements. Overall, the success of our study shows that MS Western is a versatile protein quantification tool that allows for a great flexibility in sample preparation.

We should note that the current study is the second to report a simultaneous quantitative assessment of multiple photoreceptor outer segment proteins. The previous analysis was part of a study describing the proteomes of whole and fractionated rod outer segments [10], which included the Absolute Protein Expression (APEX) approach that is based on the assumption that the frequency of peptide identification for a given protein is directly proportional to its abundance [41]. However, the authors of [10] noted that the molar ratios amongst many proteins suggested by APEX were far from independently established values. For example, the ratio between PDE6 and rhodopsin was calculated to be an order of magnitude higher than the previously reported range [42-45], while the amount of the CNG channel was about the same as PDE6 despite being known to be less abundant [46]. These and other examples in [10] illustrate that APEX may provide only rough estimates of relative protein abundances. For this reason, we did not include the APEX values from [10] in **Table 1** to avoid confusion.

### Quantification of phototransduction proteins

We will now relate our current data on outer segment protein abundances to previously published values (**Table 1**), starting with the proteins responsible for activation of the phototransduction cascade: rhodopsin, transducin (G_t_) and PDE6 (shown at the upper disc in **Fig. 1B**). We determined that the outer segment contents of the α and β subunits of G_t_ are essentially equimolar, which is consistent with these subunits being a part of a functional αβγ heterotrimer [47]. Both G_t_ subunits are present at an ∼1:8 molar ratio to rhodopsin, which matches previous measurements in mammalian rods based on semi-quantitative Western blotting [42, 48]. Similarly, our ∼1:140 ratio between the catalytic subunits of PDE6 and rhodopsin falls within the range of reported values [42-45].

Another group of proteins, shown at the middle disc in **Fig. 1B**, is involved in deactivation of the phototransduction cascade. Rhodopsin is deactivated through phosphorylation by GRK1 (*aka* rhodopsin kinase), followed by arrestin binding to phosphorylated rhodopsin; the activity of GRK1 is regulated by the Ca^2+^-binding protein, recoverin [1]. The published estimates of the molar ratio between GRK1 and rhodopsin range widely from 1:360 [13] to 1:40 [14]. We now show that this ratio is ∼1:200. The cases of recoverin and arrestin are more complex because light causes a major redistribution of these proteins between the outer segment and the cell body [49]. The reported ratio between recoverin and rhodopsin in dark-adapted mouse rod outer segments is 1:104 [50], which is confirmed by our measured ratio of 1:111. The molar ratio between arrestin and rhodopsin in isolated dark-adapted mouse rod outer segments was reported to be within the range of 1:7 to 1:10, based on semi-quantitative Western blotting [51]. However, another study analyzing the arrestin content in tangential serial sections from a frozen flat-mounted retina argued that the upper limit of arrestin content in dark-adapted outer segments is ∼7% of its total cellular amount, which corresponded to an ∼1:18 molar ratio to rhodopsin [52]. We now show that this ratio is actually equal to 1:18.

Also involved in deactivation of the phototransduction cascade is the GTPase activating complex for G_t_, which is a constitutive trimer of RGS9, Gβ5 and R9AP [37]. As discussed above, our experiments yielded an exceptional consistency in determining the outer segment abundances of these subunits averaging at an ∼1:460 molar ratio to rhodopsin. Previous estimates ranged widely between 1:270 and 1:1,640 [11, 12, 44], and the current value falls approximately in the middle of the range from the two most recent reports [12, 44].

The next two proteins, the CNG channel and the Na^+^/K^+^/Ca^2+^ exchanger (NCKX1), reside in the plasma membrane enclosing the outer segment (**Fig. 1B**). They are involved in the generation and regulation of the photoreceptor’s electrical response to light [53]. The functional CNG channel in rods is a constitutive tetramer containing three CNGα1 and one CNGβ1 subunits. Our measurements corroborated this 3:1 stoichiometry nearly perfectly and showed that the tetrameric channel is present at an ∼1:990 molar ratio to rhodopsin, which is an approximately twice lower content than estimated previously [46]. We also found that NCKX1 is present at an ∼1:470 ratio to rhodopsin, which is about twice higher than previously reported [54] and twice higher than the amount of the CNG channel. The CNG channel and NCKX1 exchanger have been shown to form a supramolecular complex in rods [55-58], and it has been suggested that two molecules of NCKX1 are bound to a single heterotetrameric CNG channel [57]. Our currently measured stoichiometry is entirely consistent with the latter model.

Two other proteins directly involved in generating the visual signal are the isoforms of guanylyl cyclase, GC1 and GC2, responsible for synthesizing cGMP, the second messenger in phototransduction. We show that they are present at ∼1:530 and ∼1:3,500 molar ratios to rhodopsin, respectively. The abundance of GC1 falls roughly in the middle of the previously reported range [8, 9, 59], while GC2 is ∼2-fold more abundant than previously reported [9, 60]. Notably these enzymes are regulated by the Ca^2+^-binding proteins, GCAP1 and GCAP2 [61], however neither GCAP1 nor GCAP2 produces a sufficient number of tryptic peptides for quantification.

### Quantification of other disc-specific proteins

Peripherin-2 and ROM1 are two homologous proteins which interact to form very large oligomeric structures that fortify photoreceptor disc rims [62] (see lower disc in **Fig. 1B**). A recent cryo-electron tomography study [63] showed that three continuous oligomeric chains of peripherin-2/ROM1 molecules span the entire circumference of a disc rim. Based on *in vitro* analyses of proteins solubilized from disc membranes, it is widely accepted that peripherin-2 and ROM1 form homo- and hetero-tetramers that are thought to serve as the base structural units of these oligomeric chains [62]. It has been reported that peripherin-2 is ∼2-2.5-fold more abundant than ROM1 [64, 65], which is well corroborated by our current data indicating that outer segments contain ∼2.0-fold more peripherin-2 than ROM1. We further showed that the molar ratio between peripherin-2/ROM1 monomers and rhodopsin is ∼1:12.7. An early estimate suggested that the presumed peripherin-2/ROM1 tetramers are present at an ∼1:90 molar ratio to rhodopsin [66]. A different value of ∼1:25 was recently calculated [67] based on fitting the dimensions of peripherin-2/ROM1 unit structures resolved by cryo-electron tomography [63] into the circumference of the disc membrane, while assuming that these units were tetramers. This would correspond to an ∼1:6.3 molar ratio between peripherin-2/ROM1 monomers and rhodopsin, which is a 2-fold higher abundance than our measured value.

However, there is a possibility to reconcile our measurement with the calculation in [67]. The cryo-electron tomography in [63], which served as the basis of that calculation, did not have sufficient resolution to determine whether the peripherin-2/ROM1 structural units observed in disc rims were dimers or tetramers, while the calculation was made with the assumption that these structures were tetramers. Yet, a recent study [68] reported a pseudo-atomic resolution model of peripherin-2/ROM1 complexes suggesting that this structural unit is a dimer. In this case, the calculation in [67] would have to be adjusted by a factor of two, which would bring it to ∼1:12.5 and match our current value of ∼1:12.7. Because our directly measured value corresponds to the model describing the base structural units of peripherin-2/ROM1 as dimers rather than tetramers, it is likely that the structural units resolved by cryo-electron tomography are actually dimers.

The last two analyzed proteins are the lipid flippases, ABCA4 and ATP8A2, uniquely expressed in discs (**Fig. 1B**, lower disc). Our measured 1:790 ratio between ABCA4 and rhodopsin is notably lower than previous measurements with mammalian rods, ranging from 1:120 [69] to 1:302 [15]. The outer segment content of ATP8A2 has only been assessed as being <0.1% of the total outer segment protein [70]. We now report that it is present at an ∼1:13,200 ratio with rhodopsin, which makes it the least abundant protein analyzed in our study.

## Concluding remarks

To better appreciate our findings, we illustrate the distribution and abundances of membrane-associated proteins quantified in this study in a cartoon representing one quarter of one lamellar surface of a photoreceptor disc (**Fig. 4**). In this illustration, each protein is represented by an object of unique shape and color, and the number of these objects corresponds to the exact number of protein molecules located in this space.

**Figure 4.**
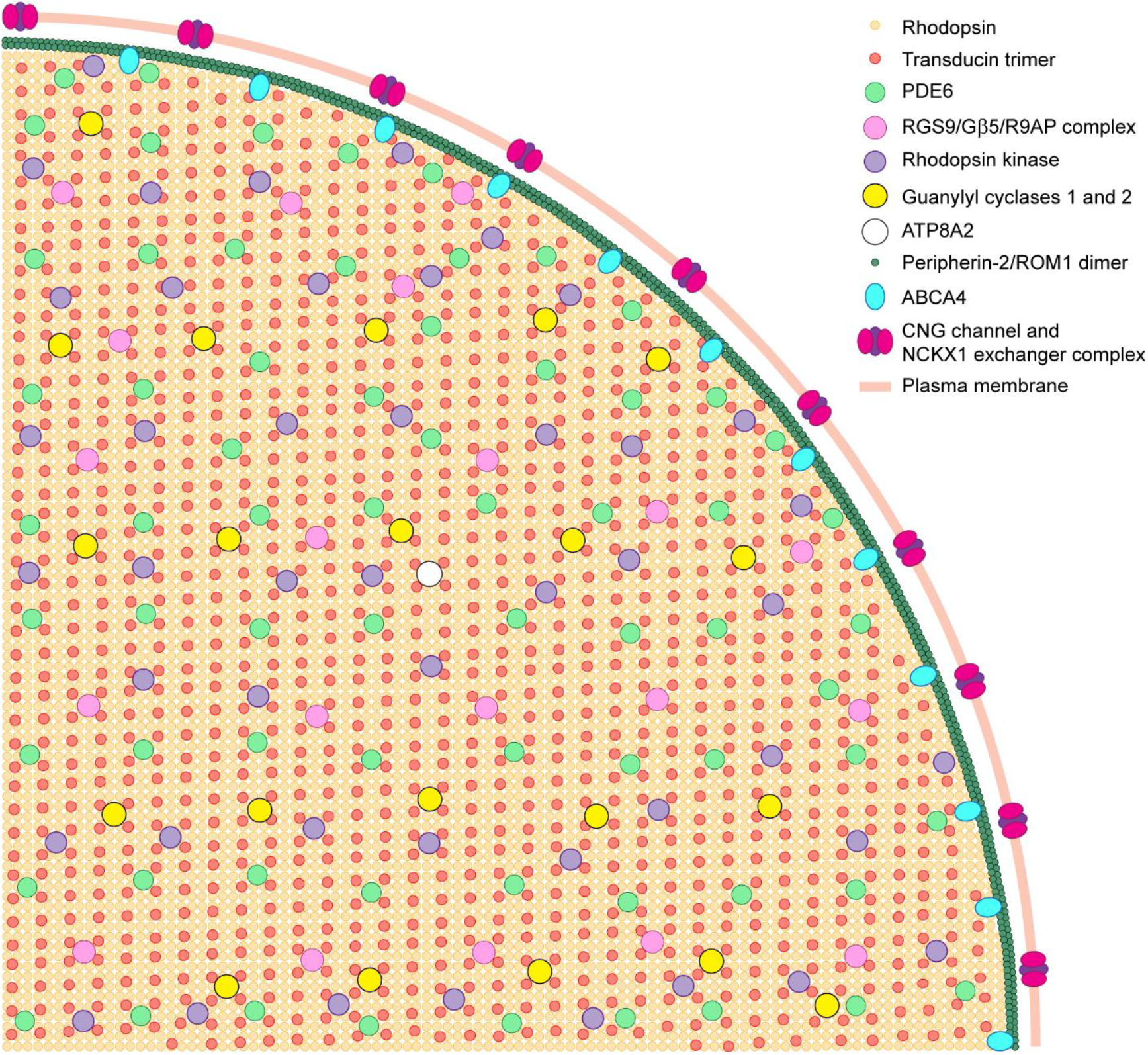
Cartoon illustrating the amounts of proteins in a section of the photoreceptor disc. The drawing shows the exact number of molecules for membrane proteins analyzed in this study within the space of one quarter of a photoreceptor disc. Shown is one disc surface and the corresponding portion of the outer segment plasma membrane surrounding this disc. The symbols representing each protein are specified in the figure. Symbol sizes do not reflect the actual dimensions of the proteins they represent; rather, less abundant proteins are shown as larger symbols for ease of visualization. Whereas plasma membrane shares many proteins with discs, only the plasma membrane specific CNG channel and NCKX1 are shown at this membrane. The peripherin-2/ROM1 structural units are shown as dimers as suggested by recent ultrastructural analyses described in the text.

One immediate observation is that the outer segment protein material is dominated by rhodopsin, which serves both as the light receptor and the essential building block of disc membranes. Our measurements indicate that rhodopsin represents an ∼92.2% molar fraction of all transmembrane proteins in the disc. The next most abundant are peripherin-2 and ROM1, which together represent ∼7.2% of the transmembrane protein fraction. The combined molar fraction of all other transmembrane proteins in disc membranes, including R9AP, GC1, GC2, ABCA4 and ATP8A2, account for only a meager ∼0.6%. Indeed, knockout of either rhodopsin or peripherin-2 completely abolishes disc formation [71, 72]. In contrast, the knockouts of R9AP, GC1, GC2, ABCA4 or ATP8A2 cause primarily functional defects in photoreceptors [73-76]. Therefore, a mature disc is essentially built from two structural protein elements: rhodopsin within the disc lamella and peripherin-2/ROM1 in the disc rim.

The second most abundant protein in the outer segment is transducin, which is peripherally associated with disc membranes. Its high concentration at the disc surface ensures a high frequency of interactions with light-activated rhodopsin, which is required for a rapid propagation of the visual signal. Another abundant outer segment protein is arrestin, which is present in an amount sufficient to deactivate the large fraction of rhodopsin activated by a bright light stimulus. The rest of the proteins analyzed in this study are expressed in catalytic amounts. Taken together, these findings contribute to our understanding of how the phototransduction pathway functions as a single, well-coordinated molecular ensemble in the spatial constraints of the outer segment architecture.

## Acknowledgements

This work was supported by the NIH grants EY030451 (VYA), EY005722 (VYA), EY033763 (TRL), the Career-Starter Research Grant from the Knights Templar Eye Foundation (WJS) and Unrestricted grant to Duke University by Research to Prevent Blindness. The authors thank Ronald Hay (University of Dundee) for providing the ΔLys, ΔArg auxotrophic strain of *E. coli*. We are grateful for Dr. Aliona Bogdanova (MPI of Molecular Cell Biology and Genetics, Dresden) for expert technical support.

## Data availability

Mass spectrometry data have been deposited to the MassIVE server at Center for Computational Mass Spectrometry University of California, San Diego, MSV000090885.

URL – ftp://massive.ucsd.edu/MSV000090885, ID – Skiban, PSW-retina5006

## Supplemental data

This article contains supplemental data.

## Supplementary Figure Legends

**Supplementary Table 1. List of peptide sequences**

Tryptic peptide sequences representing each outer segment protein of interest, which are either directly engaged in phototransduction (entries 1-16) or are specific to photoreceptor discs (entries 17-20). The LC-MS/MS sequence coverage of the chimera was 100%. Six peptides that were systematically excluded from the final analysis are indicated in red (see text for an explanation of exclusion criteria).

**Supplementary Table 2. Data used for protein quantification – Experiment 1**

**Supplementary Table 3. Data used for protein quantification – Experiment 2**

**Supplementary Table 4. Data used for protein quantification – Experiment 3**

**Supplementary Table 5**. Data **used for protein quantification – Experiment 4**

**Supplementary Table 6. Data used for protein quantification – Experiment 5**

**Supplementary Table 7. Data used for protein quantification – Experiment 6**

**Supplementary Figure 1.**
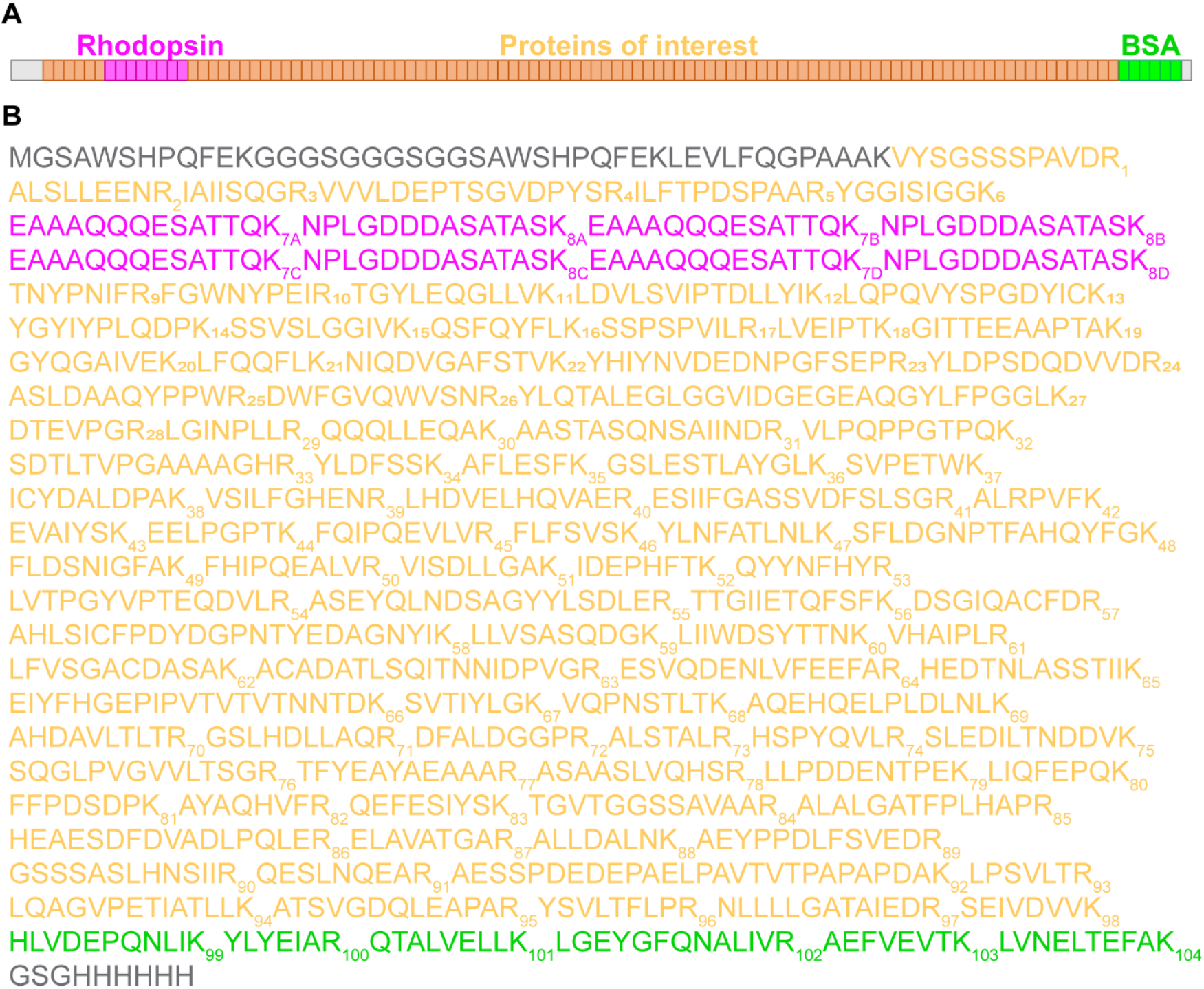
Sequence of the chimeric protein. **A**. Cartoon illustrating the composition of the chimeric protein. The chimeric protein contains concatenated sequences of 110 peptides, including 8 from rhodopsin (magenta), 96 from other outer segment proteins of interest (orange) and 6 from BSA (green). The protein also contains an N-terminal twin-strep-tag with a 3C protease cleavage site and a C-terminal His_6_-tag (grey), which were not utilized in the current study. **B**. The peptide sequence of the chimeric protein, including peptides from rhodopsin (magenta), other outer segment proteins of interest (orange) and BSA (green). The N-terminal twin-strep-tag with a 3C protease cleavage site and the C-terminal His_6_-tag are shown in grey.

